# Environmental drivers of metabolomic profiles within and between cryptic lineages of *Montastraea cavernosa*, the great star coral

**DOI:** 10.64898/2026.05.15.725494

**Authors:** Dominique N. Gallery, Evelyn A. Abbott, Lisa A. Rose Mann, Alexa K. Huzar, Karim D. Primov, Camille P. Brown, Payton L. Bryant, Brian E. Sedio, Mikhail V. Matz

## Abstract

Reef restoration practitioners aim to preserve coral genetic diversity by protecting reefs and cultivating diverse genotypes in coral nurseries. However, cryptic genetic lineages in most corals complicate restoration strategies, as the role of between-lineage genetic divergence remains unclear regarding adaptation. In *Montastraea cavernosa*, researchers have identified cryptic lineages, some strongly segregated by depth. We conducted a ten-week reciprocal transplantation experiment using two cryptic lineages restricted to shallow water (<10m depth), with one lineage more common on nearshore reefs and the other on offshore reefs. We aimed to quantify lineage-specific responses to the environment that explain the genetic and ecological divergence between the two lineages. Surprisingly, the strongest response was not lineage-specific. Instead, both lineages exhibited strong and similar changes in growth and metabolomic profiles, depending on the transplantation habitat. These results suggest that cryptic lineages employ similar mechanisms of adaptation and acclimatization to environmental challenges, despite their genetic distinction.

## Introduction

As anthropogenic stressors intensify, coral populations will continue to decline, reducing genetic diversity and, thus, corals’ ability to adapt to future conditions (Carpenter *et al*. 2008). The challenges of managing coral genetic diversity are complicated by increasing evidence for subpopulations of nominal species (Grupstra *et al*. 2024; Hays *et al*. 2021; Knowlton 1993), known as cryptic genetic lineages. Species with cryptic genetic lineages appear identical but exhibit limited introgression across the genome due to reduced interbreeding between lineages (Grupstra *et al*. 2024). Limited introgression between these lineages exacerbates genetic diversity loss due to restricted gene flow (Bálint *et al*. 2011) and makes it challenging for managers to make informed conservation decisions during restoration. Local adaptation may be one driver of this cryptic genetic structure associated with environmental variation, resulting in subpopulations with limited interbreeding in close geographic proximity (Black *et al*. 2025b; Cabacungan *et al*. 2025; Grupstra *et al*. 2024; Rippe *et al*. 2021; Sanford & Kelly 2011). While these lineages are known, we lack understanding of the environmental parameters that drive and maintain them across a seascape (Gallery *et al*. 2024; Grupstra *et al*. 2024; Rippe *et al*. 2021).

Genotype × environment (G×E) interactions occur when different genotypes respond distinctively to environmental variations, leading to differences in traits (Ottman 1996). These interactions illuminate how genetic predispositions and environmental conditions collectively influence coral phenotypes (Hackerott *et al*. 2023; Million *et al*. 2022). They can affect key traits, including growth rates, reproductive success, stress tolerance, and metabolomic profiles (Farag *et al*. 2018; Roach *et al*. 2021). Understanding the cumulative effect of these two forces is crucial to our study of resilience and adaptability in coral species’ response to changing environmental conditions (Drury *et al*. 2017). For example, certain coral genotypes may exhibit enhanced growth and survival in specific habitats and struggle in others, at least in the short term (Drury *et al*. 2017). A better awareness of GxE interactions can aid restoration efforts by guiding the selection of coral strains for outplanting in degraded reefs, ensuring that individuals are tailored to the environmental contexts of target sites.

Reciprocal transplant experiments are valuable for studying G×E interactions, particularly in clonal, benthic species such as coral (Lasky *et al*. 2023; Ondarra & Milagrosa 2020). In this study, we designed a reciprocal transplant experiment with the great star coral, *Montastraea cavernosa*, a ubiquitous, annually gonochoristic, broadcast-spawning coral found across the Atlantic Ocean, Caribbean Sea, and the Gulf of Mexico (Frys *et al*. 2020; Goodbody-Gringley *et al*. 2012). This species has multiple cryptic genetic lineages across a depth and habitat gradient in the Florida Keys (Gallery *et al*. 2024; Rippe *et al*. 2021; Sturm *et al*. 2023). Here, we used two shallow-specialized cryptic lineages, which are almost exclusively found at depths <10 meters. These lineages, referred hereafter as MC4 and MC7, were first identified by Rippe et al. (2021) and labeled “Nearshore” and “Offshore,” respectively. We renamed the lineages based on the median depth at which the individuals were found. MC4 is characterized by living predominantly in nearshore, shallow environments in the Florida Keys (Gallery *et al*. 2024; Rippe *et al*. 2021), in a habitat with high thermal variability and terrestrial nutrient influxes (Lirman & Fong 2007). MC7 lives predominantly in offshore, shallow environments in the Florida Keys (Gallery *et al*. 2024; Rippe *et al*. 2021), which have relatively fewer thermal extremes due to the proximity to the Florida Current (Kenkel & Matz 2016; Lapointe *et al*. 2004; Lee & Williams 1999) and less exposure to terrestrial pollution. Despite a preference for specific habitats, the lineages often co-occur on the same reef, making them ideal candidates for a reciprocal transplant experiment.

The literature has mixed results regarding environmental specialization, specifically regarding whether the native habitat improves the fitness of coral compared to a foreign habitat (Castillo *et al*. 2024; Clements *et al*. 2024; Drury *et al*. 2017; Marhoefer *et al*. 2021). However, because there is evidence for environmental specialization in *M. cavernosa* lineages (Rippe *et al*. 2021), we hypothesized that fragments moved to a foreign environment would experience reduced fitness relative to their clonal counterparts kept in their native environment. Further, based on work showing that the metabolomics of the coral holobiont can predict coral genotypes (Williams 2024), we predicted that chemical composition would be most strongly influenced by lineage assignment.

## Methods

### Initial sampling

One hundred and sixty tissue samples of *Montastraea cavernosa* were collected and tagged from three locations in the Florida Keys — Carly’s Patch (24.6071, -81.4294), Haslun’s Heads (24.5532, -81.4376), and Looe Deep Reef (24.5410, -81.4141) (Figure 1a). Tissue was collected using a 20 mL needleless syringe to isolate a single polyp per colony. Following tissue collection, the syringes were placed on ice and transported to Mote Marine Laboratory’s Elizabeth Moore International Center for Coral Reef Research & Restoration (IC2R3). Sea water was removed from the sample using a glass fiber prefilter (Merck Millepora Ref. No. AP2501300), and the remaining polyp was placed in 100% ethanol and stored in a -80°C freezer until transportation to the University of Texas at Austin. Samples were transported on dry ice and stored at -80°C until processed.

**Figure 1:**
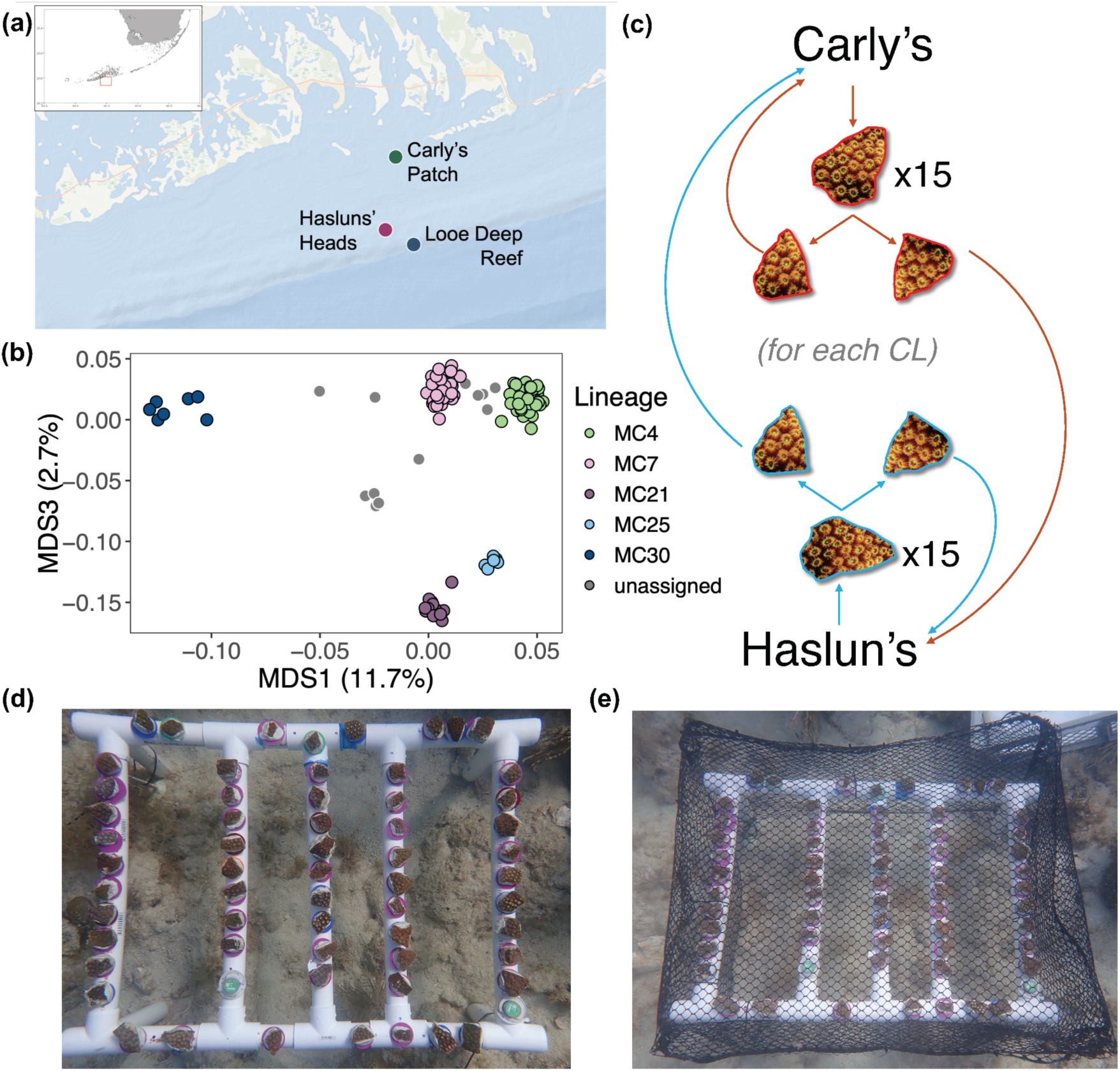
(a) Location from which samples were collected and tagged. Samples for this study were collected at Carly’s Patch and Haslun’s Heads. (b) Principal coordinate analysis of genetic variation among samples collected for this study and previously published samples (Gallery *et al*. 2024; Rippe *et al*. 2021). Samples belonging to the MC4 and MC7 lineage found at Carly’s Patch and Haslun’s Heads were selected for this study. (c) Design of the reciprocal transplant. A total of 15 samples of MC4 and 15 samples of MC7 were collected from each Carly’s Patch and Haslun’s Heads, fragmented into two pieces, and one piece of each fragment was transplanted to Carly’s Patch and Haslun’s Heads. (d) Photo of the rig at installation at Haslun’s Heads. (e) Photo of the netting used to protect the coral from herbivory at Haslun’s Heads.

### Genotyping and lineage assignment

DNA was extracted from tissue samples using a modified phenol-chloroform extraction method (https://docs.google.com/document/d/1xReYr-1CWllOhy8CnD-hHOWjFWfbfUH7/edit?usp=sharing&ouid=102775755787957987804&rtpof=true&sd=true). Extracted DNA was stored at -20°C until library preparation. The extracted DNA was cleaned using the Qiagen DNeasy PowerClean Pro Cleanup Kit (CAT No. 12997-50) following the manufacturer’s instructions. Following cleaning, samples were prepared for 2bRAD library sequencing according to the protocol at https://github.com/z0on/2bRAD_denovo. Sequencing was performed on the Illumina NovaSeq 6000 S1 platform at the University of Texas at Austin Genomic Sequencing and Analysis Facility (GSAF). Raw read sequences were processed using the methods outlined in Rippe et al. (2021) and Gallery et al. (2024). Reads were cleaned and demultiplexed following the reference-based walkthrough available at https://github.com/z0on/2bRAD_denovo/blob/master/2bRAD_README.sh. Cleaned reads were then quality-filtered using CUTADAPT version 3.5 (Martin 2011) and mapped to the *M. cavernosa* genome (Rippe et al. 2021, https://www.dropbox.com/s/yfqefzntt896xfz/Mcavernosa_genome.tgz) with bowtie2 version 2.4.5 (Langmead & Salzberg 2012). Files were converted to BAM format using SAMtools version 1.15 (Danecek *et al*. 2021; Li *et al*. 2009).

Genetic distances between samples were computed with ANGSD version 0.937 (Korneliussen *et al*. 2014) using single-read resampling identity-by-state (IBS) algorithm with the following filtering parameters: base call quality greater than Q25, mapping quality greater than 20, SNP p-value less than 1 x 10^-5^, a minimum minor allele frequency of 0.05, and sequenced in approximately 80% of samples. Reads with multiple best hits and triallelic sites were discarded. We then used the *hclust* function in R version 4.0.3 (R Core Team 2023) to create a hierarchical clustering tree to identify clones by comparing clustering to known technical replicates. One individual from each set of clones and technical replicates was retained for downstream analysis (n = 17).

After removing clones and technical replicates, we combined the samples with individuals of known lineage assignment from previous datasets (Gallery *et al*. 2024; Rippe *et al*. 2021), yielding a total of 392 individuals. We identified variant SNPs with the above ANGSD filtering parameters and retained 7346 SNPs. Moreover, we identified probable admixture assignments using NGSadmix (Skotte *et al*. 2013) for K = 2-10. We also identified the lineages of the new samples by clustering them with previously published individuals (Figure 1b). Individuals with greater than 25% assignment in a second lineage (between-lineage hybrids or representatives of undescribed rare lineages) were removed from the selection process for the reciprocal transplant. We selected MC4 lineage individuals from Carly’s Patch (n =15), MC7 individuals from Carly’s Patch (n = 15), MC4 lineage individuals from Haslun’s Heads (n = 15), and MC7 individuals from Haslun’s Heads (n = 15) (Figure 1c). Colonies were selected at random from the complete list of potential colonies.

### Reciprocal transplant

We removed approximately 10 cm^2^ of each selected colony and transplanted the fragments to Mote Marine Laboratory’s IC2R3 facility for initial processing. Each fragment was split into two pieces, attached to a bolt and tag, and placed in a common tank for 2 days. Prior to deployment on the reef, we recorded the buoyant weight of each fragment, bolt, and tag. Fragments were attached to a PVC rig (Figure 1d) and covered with a plastic mesh to protect the fragments from fish (Figure 1f). Two HOBO loggers were deployed with each rig to measure temperature at 5-minute intervals throughout the experiment. The rigs were placed at Carly’s Patch and Haslun’s Heads for ten weeks (May 23 - July 28, 2023), at which time the rigs were removed and brought to Mote Marine Laboratory’s IC2R3 facility. Fragments were removed from the rigs, and growth was measured using the buoyant weight of the fragment, bolt, and tag. Following buoyant weight measurement, we removed two polyps and stored them in 100% ethanol at -80°C for chemical analysis.

To determine whether temperatures differed significantly between the two sites during the experiment, we used R version 4.0.3 (R Core Team 2023) to calculate daily and weekly means, minimums, and maximums using data from the HOBO loggers. Statistical data were analyzed using linear mixed models with the function *lmer* using the formula mean_temp ∼ Location + (1 | DateOnly). This formula accounts for potential error introduced by repeated temperature measurements.

### Chemical extractions

Chemicals from each sample were collected using a modified version of the Sedio lab metabolomics extraction protocol (Sedio *et al*. 2018, 2021). We used an ethanol extraction method instead of a methanol-based extraction method because the samples were preserved in 100% ethanol. We removed 1.5 mL of 100% ethanol from each sample and centrifuged the ethanol for 30 minutes at 4°C at maximum speed to separate particulates from the ethanol. The supernatant was removed and placed in a vacuum centrifuge for 2 to 3 hours until approximately 0.25 mL remained. Samples were then placed in a lyophilizer for approximately 12 hours until all remaining ethanol had evaporated. The lyophilized samples were placed in a -20°C freezer until resuspension occurred. Samples were resuspended in 0.5 mL of 100% ethanol and vortexed for 5 minutes. Samples were then filtered using a 0.2-micron filter and diluted into 80% ethanol with molecular-grade water. Metabolomics data were collected using a liquid chromatography mass spectrometry method on a Thermo Scientific (Waltham, MA, USA) QExactive quadrupole-orbitrap mass spectrometer at the Biological Mass Spectrometry Facility at the University of Texas at Austin. Instrumental methods are described in Sedio et al. (2021).

### Data analysis of buoyant weight and chemical composition

Because larger coral fragments grow at a faster rate than smaller corals (Lizcano-Sandoval *et al*. 2018), we neutralized this effect by taking the residuals of the growth (final weight - initial weight) using the linear model function (*lm*) in R version 4.0.3 (R Core Team 2023) with the following formula: weight_final∼weight_inital. We then plotted the residuals of the weight, comparing genetically identical individuals across treatments using ggplot2 (Wickham 2016). We then used the function *lmerTest* (Kuznetsova *et al*. 2017) to build a linear mixed-effects model of the residuals using the following formula: resid∼Treatment+(1 | Sample_ID). Moreover, using the package emmeans (Lenth 2024), we calculated the estimated marginal means between each treatment group of the linear mixed effects model. We determined the treatment groups that had significantly different growth between pairwise samples.

To analyze the chemical differences between the treatment groups, we initially processed the samples using the program MZMine version 3.9.0 (Schmid *et al*. 2023). We quantified the structural similarity of unique compounds by constructing a feature-based molecular network (Nothias *et al*. 2020) using the Global Natural Products Social (GNPS) Molecular Networking platform (Wang *et al*. 2016). We predicted molecular formulae using SIRIUS 4 (Dührkop *et al*. 2019), predicted molecular structures using the CSI:FingerID module (Dührkop *et al*. 2015), and classified metabolites according to the NPClassifier biochemical taxonomy (Kim *et al*. 2021) using the CANOPUS (Dührkop *et al*. 2021) module of SIRIUS. Using the results from MZMINE, GNPS, and SIRIUS, we created a master data frame of predicted formulae, structures, and classifications, and the quantity of each compound in each sample for 9806 unique compounds. We then used the R package pheatmap (Kolde 2019) to plot the similarities of the chemical composition of each sample with respect to all compounds and separately for each core biosynthetic pathway represented in NPClassifier: alkaloids, amino acids and peptides, carbohydrates, fatty acids, polyketides, shikimates and phenylpropanoids, and terpenoids.

We excluded compounds with a median count of fewer than 50, leaving 2191 compounds. We normalized the remaining counts using the functions *vst* and *assay* from the R package DESeq2 (Love *et al*. 2014). Using these normalized counts, we performed a principal coordinate analysis (PCoA) with the *capscale* function in the R package vegan (Oksanen *et al*. 2022). We plotted the resultant ordinations using ggplot2 (Wickham 2016).

## Results

### Environmental monitoring

We monitored site temperatures using HOBO loggers attached to the deployed rigs at both sites. We found the mean weekly temperatures at Carly’s Patch were slightly higher than the mean weekly temperature at Haslun’s Heads (Supplementary Tables 1 and 2). This trend was also seen on the daily mean, minimum, and maximum temperatures (Figure 2), and linear mixed model results indicated a significant difference in daily mean temperature between sites (p < 2×10^-16^).

**Figure 2:**
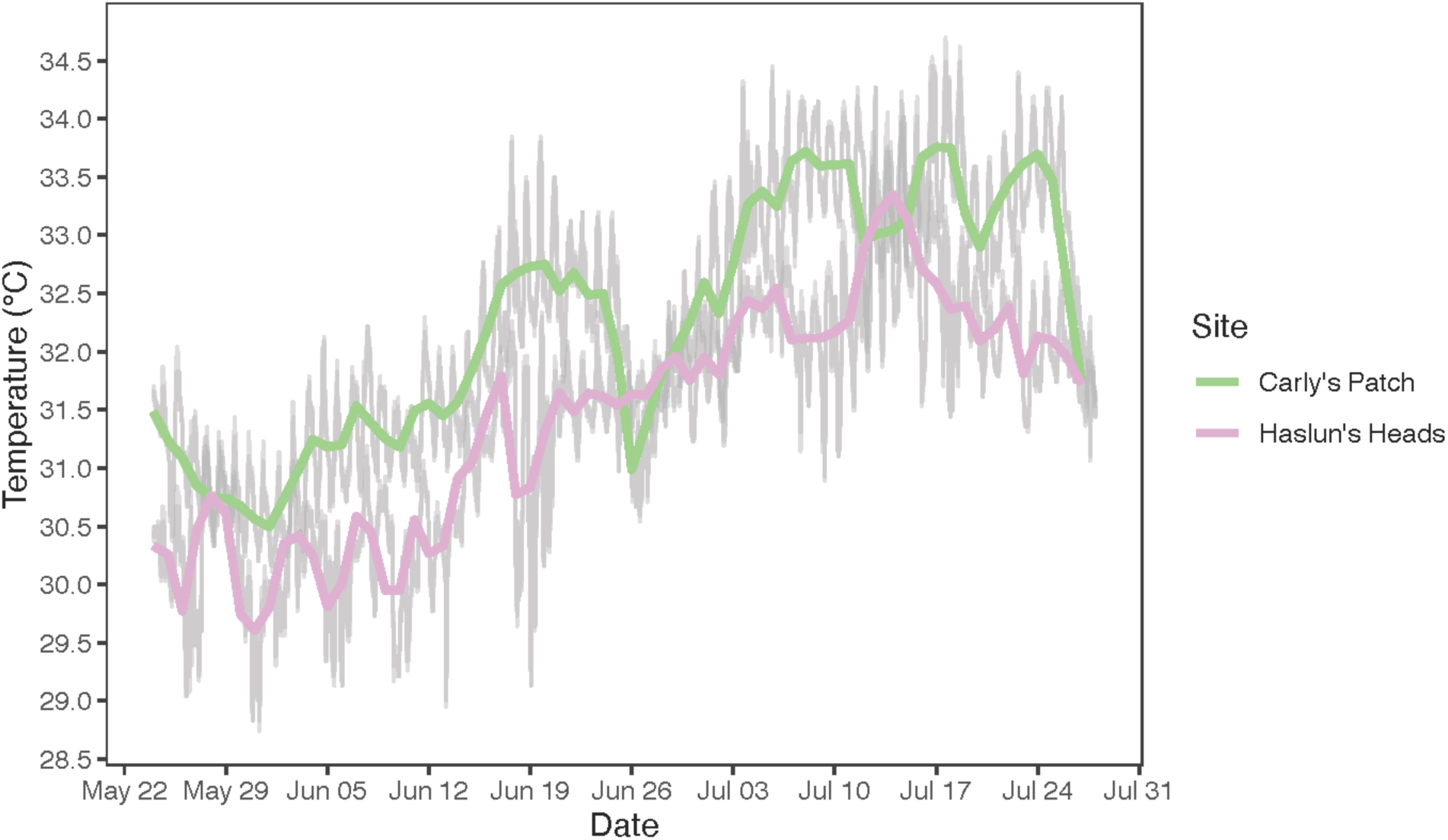
Daily maximum, minimum, and mean temperatures during the rig deployment at Carly’s Patch (green) and Haslun’s Heads (pink).

### Buoyant weights

Using the residuals from a linear model, we plotted the weight gain of each fragment and compared each to its clone in the other treatment (Figure 3). We found three groups with significant differences in weight: individuals in the MC4 lineage from Carly’s Patch, individuals in the MC7 lineage from Carly’s Patch, and individuals in the MC4 lineage from Haslun’s Heads. In these three comparisons, the mean weight gain was greater in individuals transplanted to Haslun’s Heads compared to the mean weight gain of those transplanted to Carly’s Patch. The MC7 individuals from Haslun’s Heads also exhibited this trend; however, they did not differ significantly.

**Figure 3:**
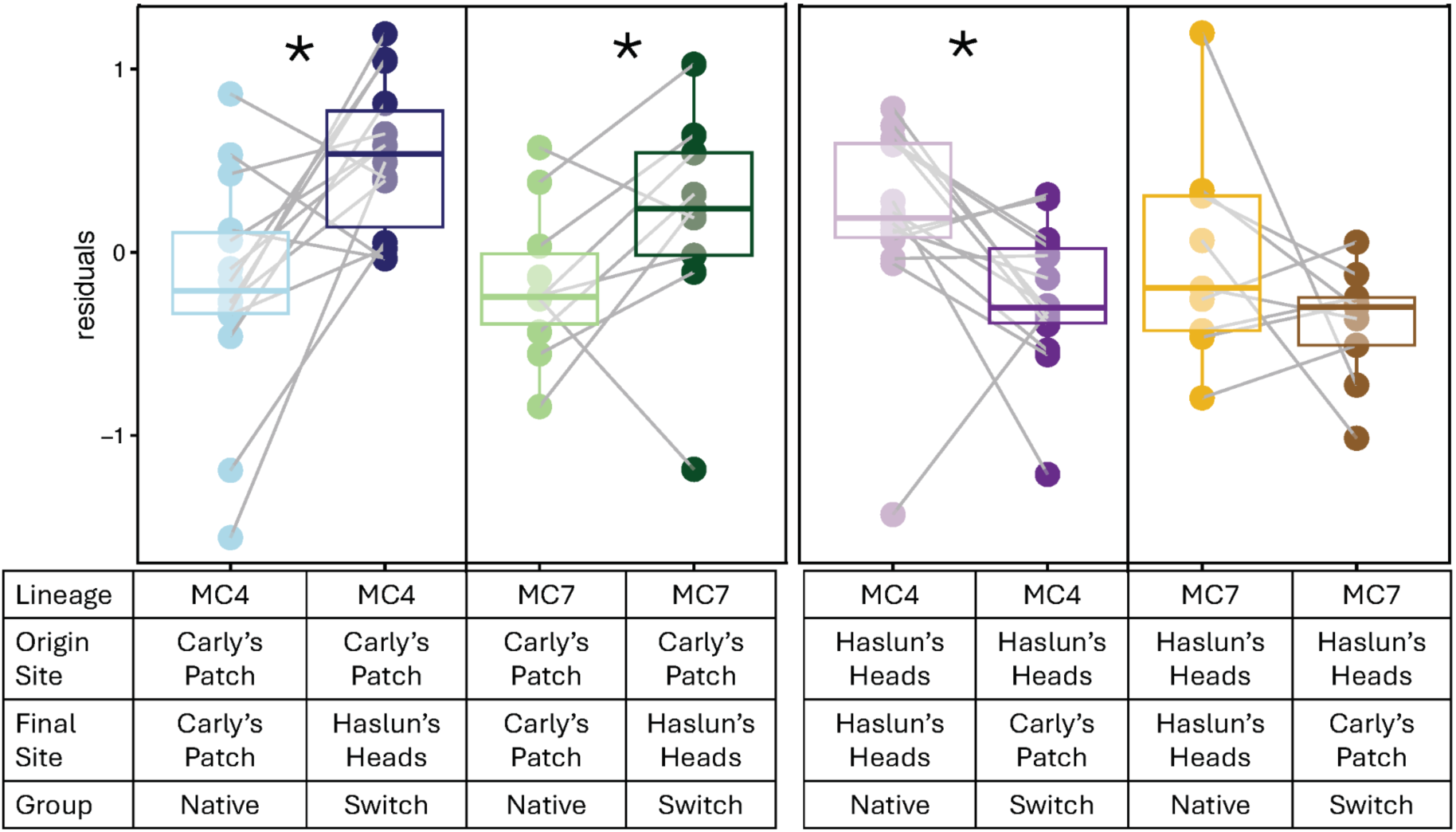
Box plot of residuals of the buoyant weight for each fragment. The light grey line connects each fragment clone. The * represents comparisons with a significant difference (p < 0.05).

### Chemical composition

We found that the chemical composition of samples was most strongly associated with their final site rather than lineage, origin site, or treatment (Figure 4). This pattern was observed across all compounds (Figure 4a), carbohydrates (Figure 4b), and amino acids and peptides (Figure 4c). The principal coordinate analysis of the chemical profiles also indicated that MDS2 is strongly associated with the final site (Figure 5), while MDS1 is associated with the lineage (Figure 5). Permutational multivariate analysis of variance (PERMANOVA) results indicated that lineage has an R^2^ = 0.09 (p ≤ 1×10^-6^), and the final site has an R^2^ = 0.24 (p ≤ 1×10^-6^). The aggregate of these results suggests that, while lineage affects coral metabolite profiles, current habitat is the strongest driver of these profiles.

**Figure 4:**
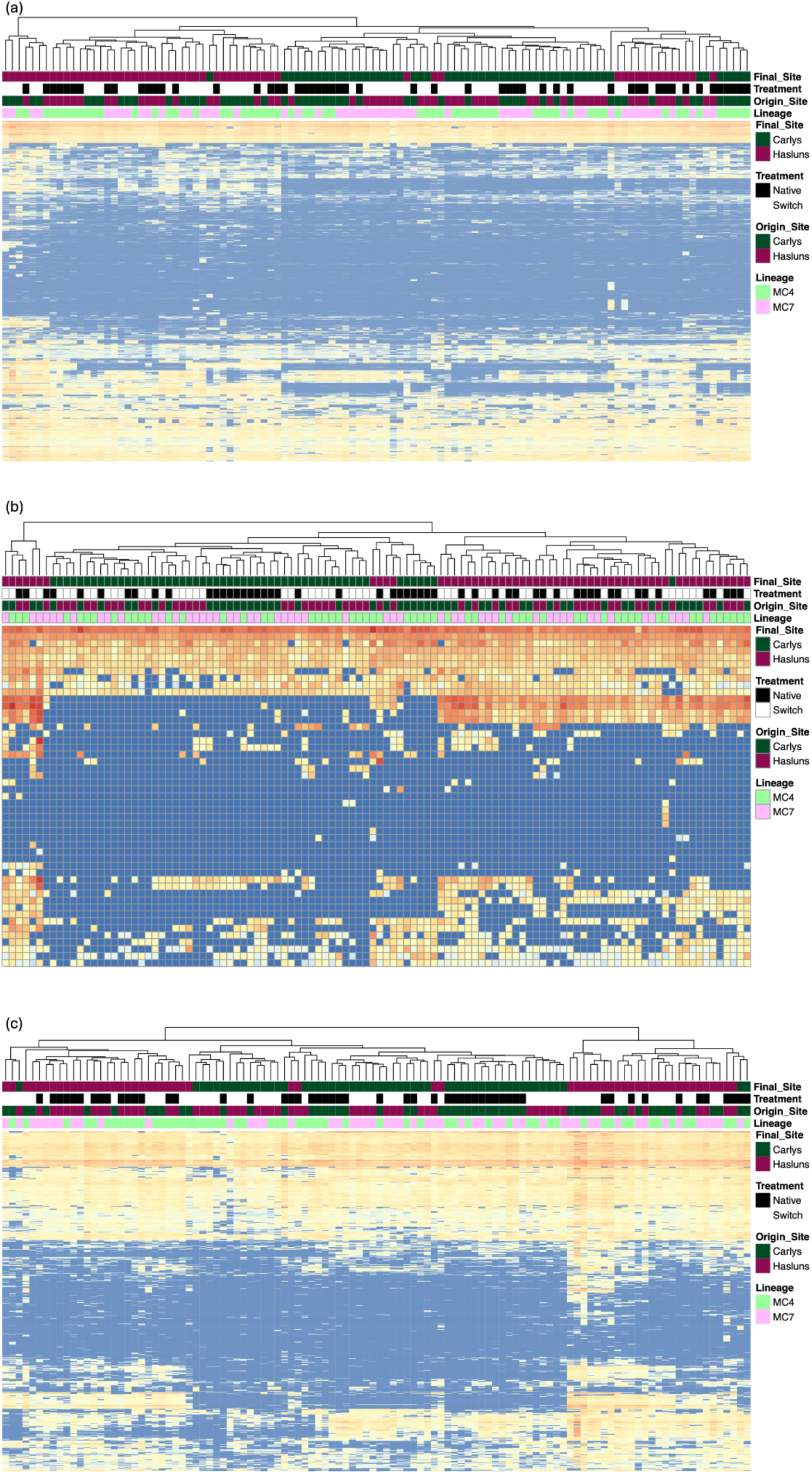
Heatmaps of the chemical composition of each sample. The x-axis represents each sample clustered by similarity, and the y-axis represents each chemical clustered by chemical similarity. (a) All compounds. (b) Carbohydrates. (c) Amino acids and peptides.

**Figure 5:**
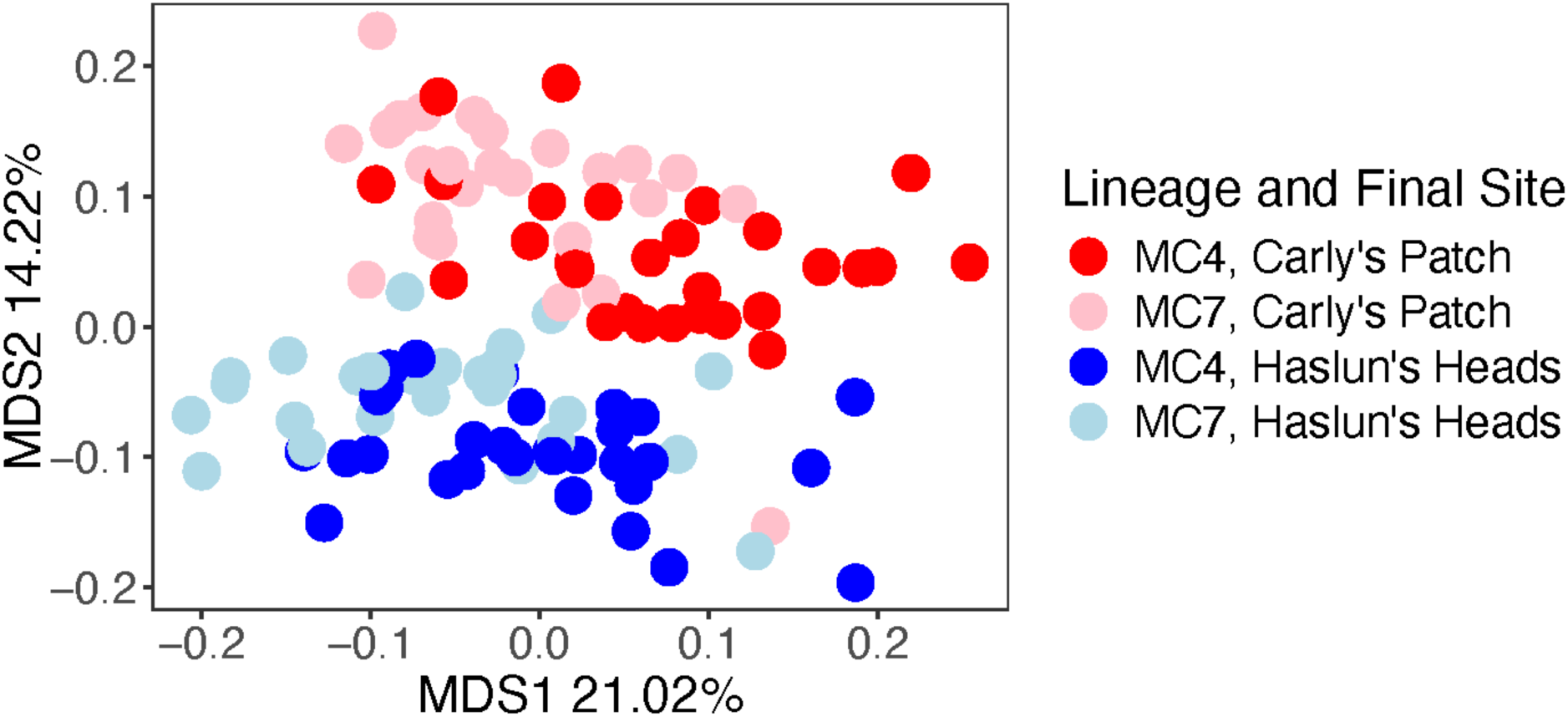
Principal component analysis of the chemical composition for all metabolites with a minimum median count greater than 50, colored by final site and shaped by lineage.

## Discussion

### Growth is predicted by the habitat of transplantation, not the lineage or habitat of origin

Our results indicate that the final habitat strongly influences corals’ growth in the MC4 and MC7 lineages of *Montastraea cavernosa*. Specifically, three of the four growth rate comparisons were significantly different, with mean growth higher in Haslun’s Heads compared to Carly’s Patch (Figure 3). The fourth comparison, although not statistically significant, showed a similar trend. This contradicts findings for the mustard hill coral (*Porites astreoides*), which showed strong trends for local acclimatization resulting in reduced fitness when transplanted to novel sites (Kenkel *et al*. 2015). In the Indo-Pacific, a year-long reciprocal transplant showed that survivorship was primarily correlated with the morphology of the coral (massive versus branching) (Tamir *et al*. 2020). Within the massive corals, physiological responses to the transplantation resulted in a loss of zooxanthellae density in individuals transplanted from deep to shallow sites (Tamir *et al*. 2020). These results counter our findings of a primarily physiological response to the transplantation site. However, a similar result was found for *Acropora millipora,* where all individuals outplanted to Keppels outperformed those outplanted to Orpheus, regardless of habitat of origin (Black *et al*. 2025a).

In the Caribbean, mangrove habitats have been suggested as potential coral refugia sites due to their ability to reduce bleaching effects by reducing light stress and temperatures (Stewart *et al*. 2021; Yates *et al*. 2014). Compared to Carly’s Patch, we suggest that Haslun’s Heads may also act as a potential refugia site for bleaching and stress tolerance due to its proximity to the Florida Current, which may reduce the average daily temperature. Moreover, our temperature profile indicated that Haslun’s Heads was cooler than Carly’s Patch at most sampling points (Figure 2). During this experiment, the Florida Keys experienced a mass bleaching event, with average daily sea surface temperatures reaching up to 31.5℃ in Key West (https://www.ncei.noaa.gov/access/coastal-water-temperature-guide/all_table.html#satl). On average, Haslun’s Heads was approximately 1℃ cooler on the PVC rig compared to Carly’s Patch. Therefore, we suggest that a longer reciprocal transplant across multiple seasons be conducted to ensure these results are not skewed by the bleaching events that now commonly occur during summer.

### Metabolomic profiles are influenced by lineage and transplantation habitat

Our results indicate a strong correlation between the transplantation habitat and lineage on the metabolomic profile of the corals (Figures 4 and 5). Understanding a coral’s metabolomics can elucidate its recent health, specifically in regard to identifying recent thermal stress (Roach *et al*. 2021; Williams *et al*. 2021). Due to an unprecedented heat wave during the experiment, the corals in this experiment experienced thermal stress; therefore, future work examining how the clones’ bleaching differs between sites would provide insight into the correlation between bleaching tolerance and the metabolome.

While recent work suggests that metabolomics can also indicate unique genotypes (Lohr *et al*. 2019), there has been less evidence to suggest that metabolomes are associated with cryptic genetic lineages. Here, we found that cryptic genetic lineages significantly differed in their metabolomic composition (Figure 4). However, these metabolomic signatures are also strongly correlated with the habitat the individuals currently reside in, which may indicate strong phenotypic plasticity of the holobiont. Since we performed chemical extractions on intact tissue, it is difficult to determine the extent to which this plasticity is attributable to the host, symbionts, or the microbiome. Furthermore, the coral microbiome plays a crucial role in coral health and response to environmental stressors (Voolstra *et al*. 2024); therefore, we suggest that future studies be performed on the gene expression of the microbiome in reciprocally transplanted coral to determine if the metabolic pathways in the microbial community are correlated with the changes we observed in the metabolome of the holobiont.

## Conclusion

Contrary to our hypothesis, individuals placed in the offshore environment grew at a faster rate than those placed in the nearshore environment, regardless of the origin site. Furthermore, while we found a significant effect of lineage on coral metabolomics, we also found that the transplantation habitat had a significant effect. These results suggest that the current habitat in which this coral species resides strongly influences its chemical phenotype and fitness. From a restoration perspective, this may indicate that the habitat in which a coral is outplanted may be more important than its cryptic lineage affiliation, at least in the short term. A longer-term experiment may be required to reveal the effect of lineage:habitat interaction on fitness.

## Acknowledgments

Funding for this project was provided by the National Science Foundation grant OCE-1737312 to M.V.M, the Burke Judd Endowment from the University of Texas at Austin to D.N.G, the Graduate Fellowship from the International Coral Reef Society to D.N.G., the AAUS Foundation Scholarship from the American Academy of Underwater Sciences to D.N.G, and Stengl Wyer Endowment grant SWG-22-01 to B.E.S. Collections and reciprocal transplant activities were authorized under the Florida Keys National Marine Sanctuaries permits FKNMS-2022-122-A1 and FKNMS-2022-122. We wish to thank the field team at Mote Marine Laboratory’s Elizabeth Moore International Center for Coral Reef Research & Restoration (IC2R3), especially Erich Bartels and Joe Kuehl, for their invaluable assistance in collecting and maintaining the project. The bioinformatics analysis was accomplished using computational resources provided by the Texas Advanced Computing Center.

## Data Accessibility

Novel sequencing data for this project will be deposited in the NCBI Short Read Archive for open access. The *Montastraea cavernosa* reference genome is available on the Matz Lab website (https://matzlab.weebly.com/data--code.html). Bioinformatic procedures associated with the de novo 2bRAD methodology (https://github.com/z0on/2bRAD_denovo) and all other data analysis procedures used in this study are available in the specified GitHub repositories (https://github.com/dgallery/MCAV_RT_git).

## Author Contributions

D.N.G., B.E.S., and M.V.M. conceived and designed this study. D.N.G, E.A.A., L.A.R.M., A.K.H., K.D.P., and C.P.B contributed to collecting samples and setting up the reciprocal transplant. D.N.G and A.K.H performed DNA extractions and 2bRAD library preparations on the samples. D.N.G. and B.E.S. performed chemical extractions on the sample. D.N.G, B.E.S., and M.V.M conducted the bioinformatic and data analysis. D.N.G., B.E.S., and M.V.M. prepared the manuscript, and all authors contributed to its final form.

## Notes

https://github.com/dgallery/MCAV_RT_git

### Competing Interest Statement

The authors have declared no competing interest.

