## Supplemental Figures for "Environmental drivers of metabolomic profiles within and between cryptic lineages of *Montastraea cavernosa*, the great star coral"

| Week | Weekly Mean | Weekly Minimum | Weekly Maximum |
| --- | --- | --- | --- |
| 5/21/23 | 31.2782477 | 27.71 | 32.6 |
| 5/28/23 | 30.7053199 | 30.11 | 31.79 |
| 6/4/23 | 31.283182 | 30.5 | 32.22 |
| 6/11/23 | 31.8090451 | 30.8 | 33.85 |
| 6/18/23 | 32.6205159 | 31.74 | 33.85 |
| 6/25/23 | 31.8580233 | 30.54 | 33.46 |
| 7/2/23 | 33.1879911 | 31.4 | 34.45 |
| 7/9/23 | 33.2807564 | 31.83 | 34.27 |
| 7/16/23 | 33.417619 | 32.34 | 34.7 |
| 7/23/23 | 31.8018576 | 27.11 | 34.32 |

Supplementary Table 1: Weekly summary temperature statistics for Carly’s Patch.

| Week | Weekly Mean | Weekly Minimum | Weekly Maximum |
| --- | --- | --- | --- |
| 5/21/23 | 30.2509747 | 27.58 | 31.27 |
| 5/28/23 | 30.18610119 | 28.74 | 31.23 |
| 6/4/23 | 30.14413938 | 29.13 | 31.19 |
| 6/11/23 | 30.91179563 | 28.95 | 32.26 |
| 6/18/23 | 31.32974206 | 29.13 | 32.34 |
| 6/25/23 | 31.75751488 | 30.97 | 32.39 |
| 7/2/23 | 32.22705853 | 31.14 | 32.9 |
| 7/9/23 | 32.730563 | 30.89 | 34.23 |
| 7/16/23 | 32.38833333 | 31.44 | 33.46 |
| 7/23/23 | 31.02020337 | 27.07 | 32.77 |

Supplementary Table 2: Weekly summary temperature statistics for Haslun’s Heads.
